# scUNVEIL: Learning scRNA-seq Cell Embeddings Through Measurement Enrichment

**DOI:** 10.64898/2026.11.17.767296

**Authors:** Tomáš Honzík, Lukáš Kuhajda, Lucie Houdová, Daniel Georgiev

## Abstract

Large-scale pretraining can improve scRNA-seq representations through *measurement enrichment* : estimating what the same measurement would reveal at greater UMI depth under the same measurement process. We introduce scUNVEIL, a self-supervised model trained through next-UMI prediction. We show that the Bayes-optimal next-UMI predictor is the posterior mean of the transcriptome composition that arbitrarily deep sequencing would reveal. Because UMI identities are exchangeable conditional on transcriptome composition, scUNVEIL operates directly on count vectors using a residual multilayer perceptron, without attention over gene tokens. The measurement-enriched transcriptome composition is predicted from a single low-dimensional hidden state, which therefore serves naturally as the cell embedding. The same model provides both reusable cell representations and measurement-enriched gene-expression estimates. scUNVEIL was pretrained without labels or auxiliary metadata on approximately 92 million human cells spanning diverse tissues. Under linear evaluation on fine-grained cell-type annotation, its zero-shot representations achieved the best performance among the evaluated methods, while providing over 1000× faster inference than the strongest evaluated transformer-based foundation model and matching that model’s predictive performance using 88% fewer input UMIs per cell. Further evaluations show that these representations improve prediction of tumor reactivity, mitochondrial membrane potential, cell morphology, and lineage-associated treatment outcomes.

## 1 Introduction

The central promise of foundation models is that broad, large-scale pretraining can produce representations that transfer across tasks and outperform specialized models trained from scratch, particularly when task-specific labels are scarce. Although remarkably successful in fields like natural language processing or computer vision, this promise remains far from fully realized in single-cell RNA sequencing (scRNA-seq). Large transformer-based models such as scGPT [5], scFoundation [12], Geneformer [26], and UCE [22] are expensive to train and deploy, while independent evaluations have found their representations to be matched or outperformed by substantially simpler baseline methods on simple benchmarks like cell-type annotation [17, 3, 25]. These results motivate a closer examination not only of architecture and scale, but of pretraining itself: what, specifically, should pretraining do to produce representations that transfer effectively across biological tasks?

Suppose an experiment measures a population of cells, yielding approximately 1,000 UMIs per cell, and a second dataset is created from the original measurement by discarding 90 % of the observed UMIs at random, yielding approximately 100 UMIs per cell. For almost any downstream analysis, the higher-depth original measurement would be preferable to work with: it contains more information about the cells’ transcriptomes, even though it may preserve, or reveal more clearly, technical distortions. Greater depth reduces uncertainty about underlying biological states, even though those states remain observed through a distorted measurement process. We call the inverse process of this random subsampling *measurement enrichment* : estimating the transcriptome composition that would be observed at greater UMI depth under the same effective measurement process. This objective does not remove systematic measurement effects. Batch effects, protocol-specific biases, and other distortions remain part of the target insofar as they shape which UMIs the assay would capture at greater depth.

scUNVEIL operationalizes this idea through next-UMI prediction. During pretraining, observed UMIs from each cell are randomly partitioned into retained inputs and held-out targets, and the model predicts the gene-identity distribution of the held-out molecules from those retained as input.

Because UMI identities are exchangeable conditional on this composition, their ordering contains no information relevant to next-UMI prediction. scUNVEIL therefore represents each cell as a fixed-length gene-count vector processed by a residual multilayer perceptron, without attention across thousands of gene tokens per cell. The architecture thus follows from the statistical structure of the objective while making representation extraction computationally efficient.

The predicted measurement-enriched transcriptome composition is decoded from a single low-dimensional hidden state. Because all information used for next-UMI prediction passes through this bottleneck, the state must retain the features most informative about the cell’s effective measured composition and therefore serves naturally as a cell embedding. The same forward pass produces two complementary outputs: a reusable cell embedding and a measurement-enriched estimate of gene-expression composition. The embedding can be transferred to downstream tasks, while the predicted composition can be used directly for gene-expression imputation.

We pretrained scUNVEIL without labels or auxiliary metadata on approximately 92 million human cells spanning diverse tissues. Under linear evaluation on fine-grained cell-type annotation, the resulting zero-shot representations achieve the best performance among the evaluated methods while requiring substantially less inference compute than transformer-based single-cell foundation models. We further evaluated the representations’ broader biological utility through supplementary experiments predicting tumor reactivity, mitochondrial membrane potential, cell morphology, and lineage-associated treatment outcomes.

## 2 Related Work

### 2.1 Denoising and count-based representations

Count-based generative models established the link between expression reconstruction and low-dimensional cell representations. scVI [19] and DCA [10] learn neural representations through count-distribution likelihoods, while SAVER-X [28] transfers information from pretrained autoencoders to improve expression estimates in new datasets. scETM [31] combines a fully connected encoder with a linear decoder and a categorical expression likelihood, and demonstrates transfer to unseen tissues and species. These methods provide important precedents for generative cell embeddings and linear expression decoding.

Molecule-level self-supervision also predates large-scale foundation models. Noise2Self [1] and molecular cross-validation [2] exploit random partitions of detected molecules to learn or assess denoisers without clean targets. Separating measurement noise from expression variation provides a related statistical foundation [23]. scUNVEIL builds on these ideas by using complementary UMI splits for categorical next-molecule prediction and reusable representation learning.

### 2.2 Large-scale cell representation learning

Large-scale self-supervision is not restricted to transformers. Richter et al. [21] studied fully connected networks pretrained on over 20M cells using masked-autoencoder and contrastive objectives, evaluating transfer across annotation, expression reconstruction, cross-modality prediction, and integration. Other MLP-based approaches use labels: SCimilarity [14] combines ontology-guided triplet learning with reconstruction, while scTab [11] scales supervised cell-type classification with a TabNet architecture. These studies establish both supervised and self-supervised precedents for scalable cell-vector representations.

Transformer-based models instead process gene-level tokens using several encodings and objectives. scBERT [29] reconstructs masked expression bins; Geneformer [26] predicts masked genes in an expression-ranked representation; tGPT [24] uses autoregressive prediction over ranked genes; and scGPT [5] learns through generative expression prediction. UCE [22] and scPRINT [16] incorporate protein-language-model gene embeddings, while TranscriptFormer [20] uses related gene features for generative modeling across 12 species. CellFM [30] explores a large RetNet-style architecture. These approaches differ in their use of ordering, biological priors, and prediction targets; gene-token attention does not itself require a meaningful molecule sequence.

A particularly relevant objective is scFoundation’s read-depth-aware pretraining [12], also used by AIDO.Cell [15]. It predicts higher-depth expression from downsampled inputs using explicit source and target depth information and masked regression on log-counts. Reconstruction-trained cell embeddings also occur within transformer models: UCE predicts masked expression status from a cell embedding and gene features, while scPRINT explicitly reconstructs expression through a cell-embedding bottleneck. scUNVEIL combines a full-transcriptome categorical next-UMI objective with a residual MLP operating on a single count vector. All next-UMI predictions pass through one cell embedding and a tied linear decoder for gene logits, without labels, auxiliary metadata, or explicit target-depth conditioning.

### 2.3 Evaluation and simple baselines

Independent evaluations show that the benefits of pretraining depend on the task and evaluation setting. Kedzierska et al. [17] found limitations of zero-shot scGPT and Geneformer representations relative to established integration and expression baselines. Boiarsky et al. [3] showed that regularized logistic regression can match or exceed scBERT on cell-type annotation, while Souza and Mehta [25] report competitive results from normalization and simple linear pipelines. GenePT’s weighted variant [4] offers another simple alternative, constructing cell vectors by expression-weighted averaging of text-derived gene embeddings. DenAdel et al. [9] found that current pretraining approaches often plateau before using their full training corpora, and De Waele et al. [8] demonstrated that input encoding, reconstruction losses, and pretraining objectives materially affect downstream performance. Together, these findings motivate evaluating the objective, architecture, and computational cost alongside strong simple baselines.

## 3 Methods

### 3.1 Training Data

We pretrained scUNVEIL on approximately 92 million human cells obtained from the CELLxGENE Census (release 2025-11-08) [6]. We included all *Homo sapiens* cells with `is_primary_data = True`. We further filtered cells with 300 or fewer detected genes and cells with 40,000 or more UMIs.

The complete list of source datasets used for pretraining is provided in Supplementary Table `Supplementary_Table_S1_training_datasets.csv`.

To avoid potential data leakage in downstream benchmarks, we explicitly excluded the Hao et al. dataset [13] from the pretraining corpus. All reported results on this dataset therefore reflect true zero-shot generalization.

The resulting corpus spans a broad range of tissues, cell types, disease states, and experimental conditions, covering 60,000 genes across the human transcriptome. Throughout this work, the term “genes” refers to the scRNA-seq features present in the reference annotation, including protein-coding genes, non-coding RNAs, and related gene-associated features.

Importantly, the model was trained using only the raw UMI count vector of each individual cell. No donor metadata, dataset identifiers, tissue labels, cell-type annotations, or any other auxiliary information was used during pretraining.

### 3.2 Pretraining Objective

We formulate pretraining as next-UMI prediction and implement this objective efficiently with an MLP backbone. We represent each cell as a single fixed-length count vector: a bag-of-RNAs representation. This allows the entire transcriptome to be processed in one forward pass, without attention over gene tokens.

For each training cell, we perform binomial subsampling with retention probability *r* ~ Uniform(0, 0.9). Restricting the upper bound reduces the frequency of splits with very small or empty held-out sets and thereby improves optimization efficiency. This randomly splits the observed UMIs into two sets: an input set and a held-out set. In the theoretical derivation, we use the idealized full-range formulation *r* ~ Uniform(0, 1) and explicitly condition on a nonempty held-out set. Operationally, the task can be viewed as predicting the identity of one randomly sampled held-out UMI from the subsampled input. Instead of sampling a single held-out UMI at each training step, however, we use the empirical gene-frequency distribution of all held-out UMIs as a soft target. For categorical cross-entropy, this is mathematically equivalent in expectation to single-UMI prediction, but is much more sample efficient.

We make this statement rigorous in Appendix A and use it to interpret the model’s output as an estimate of the measurement that would be obtained at arbitrarily deep sequencing.

### 3.3 Measurement enrichment and next-UMI prediction

For cell *c*, let **p**_c_ *∈* Δ^G*−*1^ denote its latent biological transcriptome composition. Biological and technical factors affect how this molecular population is converted into captured UMIs. We represent their combined action by an effective observation channel 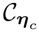 and define

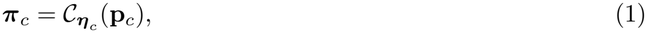

where ***π***_c_ *∈* Δ^G*−*1^ is the cell’s *effective measured transcriptome composition*. Its component *π*_c,g_ is the probability that an additional UMI generated under the same effective measurement process is assigned to gene *g*.

The resulting observation chain is

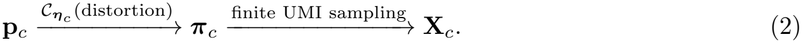

Conditional on the observed UMI depth *N*_c_, we use the idealized model

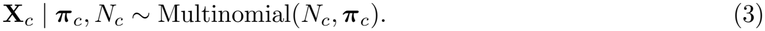

scUNVEIL addresses the second arrow in Equation (2): it estimates ***π***_c_ from the finite observation **X**_c_. It does not attempt to invert the observation channel or directly recover **p**_c_.

#### Training objective

During training, the observed UMIs of each cell are randomly divided into retained input counts and held-out target counts,

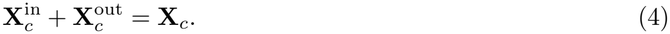

Conditional on ***π***_c_ and the split sizes, both vectors are independent multinomial samples from the same effective composition:

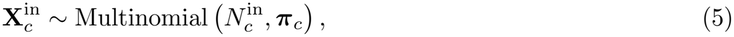

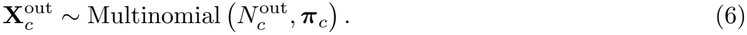

Splits with 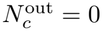 are excluded. Defining the normalized held-out composition as

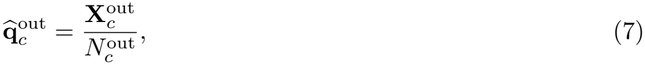

the model is trained using

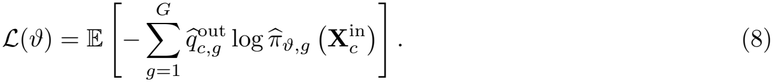

Under categorical cross-entropy, the Bayes-optimal predictor is

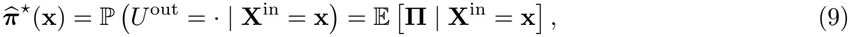

where **Π** denotes the population distribution of effective measured compositions and *U*^out^ is a uniformly selected held-out UMI. Thus, next-UMI prediction estimates both the posterior predictive distribution of an additional UMI and the posterior mean of the effective measured composition. The complete derivation, including the relationship between the single-UMI and empirical held-out objectives, is provided in Appendix A.

#### Model output and interpretation

Let **z**_θ_(**x**) *∈* ℝ^d^ denote the output of the residual MLP before its final prediction layer. The model predicts

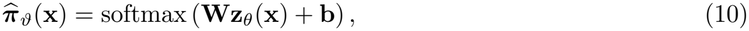

where **z**_θ_(**x**) is used as the scUNVEIL cell embedding. We define the learned measurement-enrichment operator as

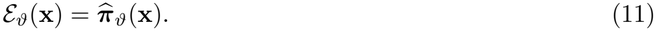

For visualization and downstream analysis, the predicted composition may be expressed as log counts per million:

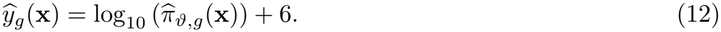

This is a rescaling of the predicted probability distribution rather than a prediction conditioned on a target depth of one million UMIs.

Measurement enrichment is an inference procedure, not the acquisition of an additional physical measurement. Cell-specific features that are absent from the observed UMIs and not predictable from the pretraining population cannot be recovered.

### 3.4 Model Architecture and Pretraining Hyperparameters

scUNVEIL uses a simple MLP backbone with residual connections and layer normalization operating on a single cell-level count vector. The input and output vocabularies consist of 60,000 CELLxGENE features, covering nearly the full human gene set represented in the Census.

The model has an embedding dimension of 2,048 and consists of 16 residual MLP blocks. Each block contains layer normalization, two dense layers of width 2,048, and an intervening ReLU activation. The final output layer maps the 2,048-dimensional representation to a probability distribution over the full gene vocabulary. As an architectural ablation, we also trained a “Linear Embeddings” variant with all residual blocks removed. Its learned transformation consists only of the input projection to 2,048 dimensions, final layer normalization, and the output projection back to the gene vocabulary, followed by softmax.

We pretrained the model for 640,000 optimization steps with a batch size of 1,000 cells, corresponding to approximately 7 full epochs over the 92-million-cell training dataset. Optimization used Adam with a cosine learning-rate schedule, starting at 10^*−*4^ and reaching 3 × 10^*−*5^ at step 640,000, with linear warmup during the first 10,000 steps.

A throughput estimate from a static batch suggested that the same optimization workload would require approximately 3 GPU-hours on an NVIDIA H100 under an optimized input pipeline. Our actual run was dominated by data-loading overhead and took approximately 70 hours on a single NVIDIA RTX A4000 GPU.

### 3.5 Embeddings PCA

After pretraining is completed, we compute a PCA projection of the embedding space. To do so, we first generate embeddings for 1,000,000 randomly sampled cells from the pretraining corpus and then fit PCA on the resulting representations. The PCA projection is provided for scenarios where a lower-dimensional representation is desired. The resulting PCA-transformed embeddings are ordered according to the significance of individual components, allowing users to select the top-*K* components based on their requirements.

## 4 Results

### 4.1 Zero-Shot Cell-Type Annotation

#### Benchmark and evaluation rationale

We first asked whether scUNVEIL embeddings make biologically relevant cell-identity information accessible to a simple linear readout. Fine-grained cell-type annotation provides an interpretable test of representation quality: the representation must separate closely related cell populations, while the limited capacity of the downstream classifier reduces the extent to which weak representations can be compensated for during task-specific training.

We evaluated classification of 58 peripheral blood mononuclear cell (PBMC) subtypes in the Hao et al. 2021 multimodal reference dataset [13]. The dataset contains 161,764 cells from 8 donors, jointly profiled using scRNA-seq and a 228-protein CITE-seq panel. Its annotations were derived from the integrated RNA and surface-protein measurements, providing high-confidence labels at substantially finer granularity than conventional major-lineage PBMC annotations. As described in the Methods, the Hao et al. dataset was explicitly excluded from the scUNVEIL pretraining corpus.

#### Evaluation protocol

For each frozen representation, we trained a linear classifier using Adam, a batch size of 1024, zero initialization, and early stopping based on a validation fold. Because representations can differ substantially in scale and optimization geometry, the learning rate was selected independently for each method from a logarithmically spaced grid with half-decade resolution:

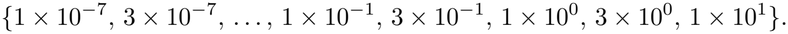

Learning rates were calibrated on a fixed four-donor training/four-donor validation split, selecting the lowest observed validation cross-entropy. Uncompetitive sweep runs were pruned using extrapolated recent validation-loss trends.

To measure generalization across individuals, we partitioned the 8 donors into 4 training donors, 3 validation donors, and 1 test donor. We repeated this procedure for 100 random donor splits per representation and report the mean ± standard deviation of macro F1, balanced accuracy, and overall accuracy.

We also measured the computational cost of generating each representation. All pipelines were run on the same AMD Ryzen 7 7700 CPU, and wall-clock time was measured for the complete 161,764-cell dataset. For dataset-specific methods that require fitting on the benchmark dataset, including scVI [19], Harmony with PCA, and MAGIC, the reported runtime includes this fitting stage.

#### Comparison design

The comparison was organized to address three complementary questions. First, we tested whether scUNVEIL improves over direct use of the observed measurements. These controls comprised the full raw 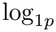-transformed expression vector and a restricted representation containing 50 manually selected PBMC marker genes listed in Table 2.

Second, we compared scUNVEIL with lightweight methods fitted directly to the benchmark dataset. Harmony with PCA and scVI were evaluated at multiple target dimensionalities. We also evaluated gene-expression imputation using MAGIC and scVI: each method was applied to the complete transcriptome, after which the same 50 marker genes were supplied to the linear classifier.

Third, we compared scUNVEIL with pretrained single-cell representation models. This is the most direct comparison because these methods, like scUNVEIL, produce reusable representations from a pretrained model rather than learning a new representation specifically for the Hao et al. dataset. Competing representations were generated using public checkpoints and recommended inference pipelines where available. Exact preprocessing and inference scripts are provided in the reproducibility repository. We selected influential pretrained models spanning different architectures and pretraining objectives.

Setting up most competing models required substantial implementation effort, and generating their representations was computationally expensive, limiting the number of methods we could evaluate. All evaluated models are included in Table 1, without exclusions based on benchmark performance.

**Table 1:**
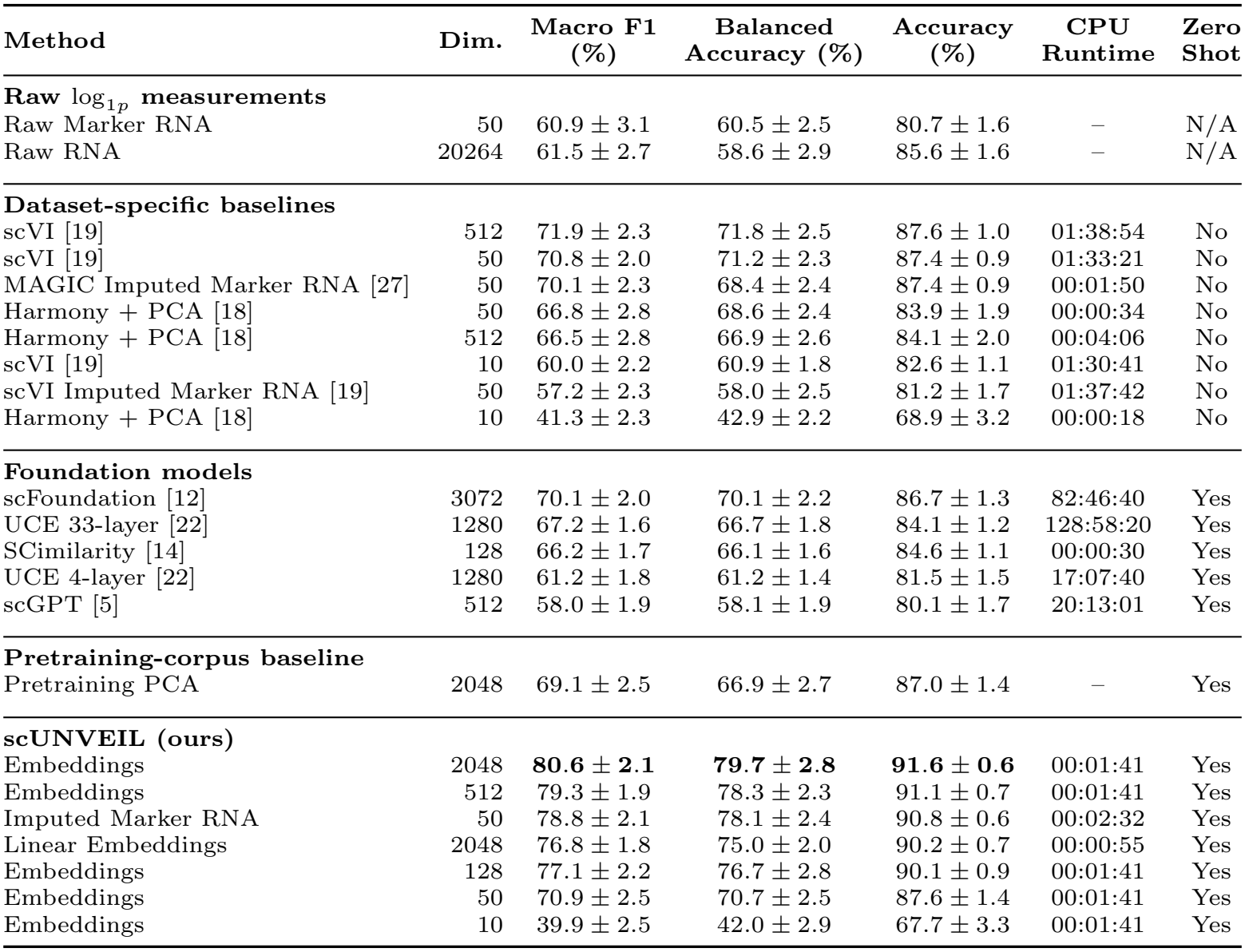
Cell-type annotation performance on the Hao et al. 2021 PBMC reference dataset, comprising 161,764 cells from 8 donors and 58 cell-type classes. Values are reported as mean ± standard deviation using 100 donor-level train–validation–test splits. The “Zero-shot” column indicates whether the representation was generated without fitting or otherwise optimizing the representation model on the Hao et al. dataset. CPU runtime is the wall-clock time required to generate representations for the complete dataset on a single AMD Ryzen 7 7700 CPU; for dataset-specific methods, this includes the time required to fit the representation model. The best result for each annotation metric is shown in bold.

| **Method** | **Dim.** | **Macro F1<br>(%)** | **Balanced<br>Accuracy (%)** | **Accuracy<br>(%)** | **CPU<br>Runtime** | **Zero<br>Shot** |
|---|---|---|---|---|---|---|
| **Raw log<sub>1p</sub> measurements** |  |  |  |  |  |  |
| Raw Marker RNA | 50 | 60.9 ± 3.1 | 60.5 ± 2.5 | 80.7 ± 1.6 | – | N/A |
| Raw RNA | 20264 | 61.5 ± 2.7 | 58.6 ± 2.9 | 85.6 ± 1.6 | – | N/A |
| **Dataset-specific baselines** |  |  |  |  |  |  |
| scVI [19] | 512 | 71.9 ± 2.3 | 71.8 ± 2.5 | 87.6 ± 1.0 | 01:38:54 | No |
| scVI [19] | 50 | 70.8 ± 2.0 | 71.2 ± 2.3 | 87.4 ± 0.9 | 01:33:21 | No |
| MAGIC Imputed Marker RNA [27] | 50 | 70.1 ± 2.3 | 68.4 ± 2.4 | 87.4 ± 0.9 | 00:01:50 | No |
| Harmony + PCA [18] | 50 | 66.8 ± 2.8 | 68.6 ± 2.4 | 83.9 ± 1.9 | 00:00:34 | No |
| Harmony + PCA [18] | 512 | 66.5 ± 2.8 | 66.9 ± 2.6 | 84.1 ± 2.0 | 00:04:06 | No |
| scVI [19] | 10 | 60.0 ± 2.2 | 60.9 ± 1.8 | 82.6 ± 1.1 | 01:30:41 | No |
| scVI Imputed Marker RNA [19] | 50 | 57.2 ± 2.3 | 58.0 ± 2.5 | 81.2 ± 1.7 | 01:37:42 | No |
| Harmony + PCA [18] | 10 | 41.3 ± 2.3 | 42.9 ± 2.2 | 68.9 ± 3.2 | 00:00:18 | No |
| **Foundation models** |  |  |  |  |  |  |
| scFoundation [12] | 3072 | 70.1 ± 2.0 | 70.1 ± 2.2 | 86.7 ± 1.3 | 82:46:40 | Yes |
| UCE 33-layer [22] | 1280 | 67.2 ± 1.6 | 66.7 ± 1.8 | 84.1 ± 1.2 | 128:58:20 | Yes |
| SCimilarity [14] | 128 | 66.2 ± 1.7 | 66.1 ± 1.6 | 84.6 ± 1.1 | 00:00:30 | Yes |
| UCE 4-layer [22] | 1280 | 61.2 ± 1.8 | 61.2 ± 1.4 | 81.5 ± 1.5 | 17:07:40 | Yes |
| scGPT [5] | 512 | 58.0 ± 1.9 | 58.1 ± 1.9 | 80.1 ± 1.7 | 20:13:01 | Yes |
| **Pretraining-corpus baseline** |  |  |  |  |  |  |
| Pretraining PCA | 2048 | 69.1 ± 2.5 | 66.9 ± 2.7 | 87.0 ± 1.4 | – | Yes |
| **scUNVEIL (ours)** |  |  |  |  |  |  |
| Embeddings | 2048 | **80.6 ± 2.1** | **79.7 ± 2.8** | **91.6 ± 0.6** | 00:01:41 | Yes |
| Embeddings | 512 | 79.3 ± 1.9 | 78.3 ± 2.3 | 91.1 ± 0.7 | 00:01:41 | Yes |
| Imputed Marker RNA | 50 | 78.8 ± 2.1 | 78.1 ± 2.4 | 90.8 ± 0.6 | 00:02:32 | Yes |
| Linear Embeddings | 2048 | 76.8 ± 1.8 | 75.0 ± 2.0 | 90.2 ± 0.7 | 00:00:55 | Yes |
| Embeddings | 128 | 77.1 ± 2.2 | 76.7 ± 2.8 | 90.1 ± 0.9 | 00:01:41 | Yes |
| Embeddings | 50 | 70.9 ± 2.5 | 70.7 ± 2.5 | 87.6 ± 1.4 | 00:01:41 | Yes |
| Embeddings | 10 | 39.9 ± 2.5 | 42.0 ± 2.9 | 67.7 ± 3.3 | 00:01:41 | Yes |

**Table 2:**
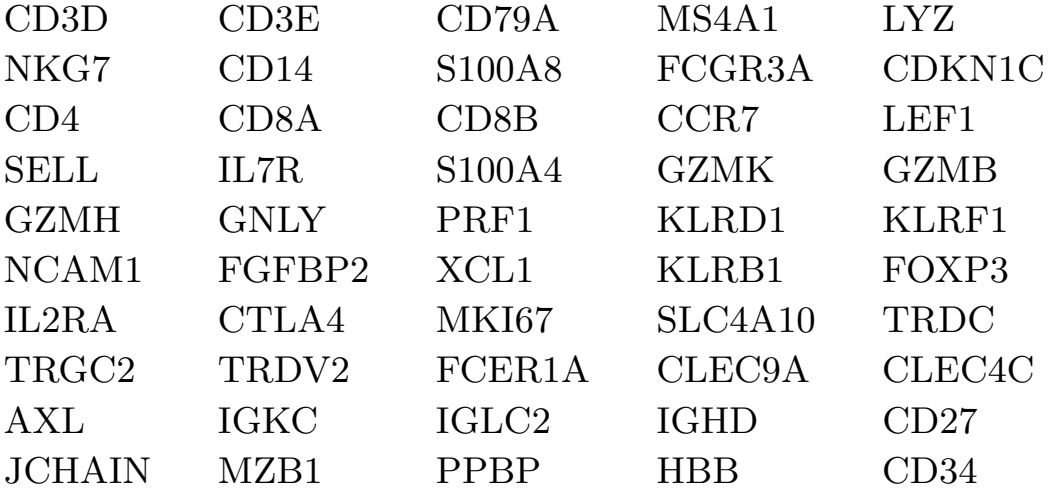
Selected 50 PBMC marker genes.

|  |  |  |  |  |
|---|---|---|---|---|
| CD3D | CD3E | CD79A | MS4A1 | LYZ |
| NKG7 | CD14 | S100A8 | FCGR3A | CDKN1C |
| CD4 | CD8A | CD8B | CCR7 | LEF1 |
| SELL | IL7R | S100A4 | GZMK | GZMB |
| GZMH | GNLY | PRF1 | KLRD1 | KLRF1 |
| NCAM1 | FGFBP2 | XCL1 | KLRB1 | FOXP3 |
| IL2RA | CTLA4 | MKI67 | SLC4A10 | TRDC |
| TRGC2 | TRDV2 | FCER1A | CLEC9A | CLEC4C |
| AXL | IGKC | IGLC2 | IGHD | CD27 |
| JCHAIN | MZB1 | PPBP | HBB | CD34 |

#### scUNVEIL produces the strongest linearly accessible representations

The full 2048-dimensional scUNVEIL embeddings achieved the highest performance on all three annotation metrics (Table 1), reaching 80.6 ± 2.1% macro F1, 79.7 ± 2.8% balanced accuracy, and 91.6 ± 0.6% overall accuracy.

Among methods fitted directly to the benchmark dataset, the strongest result was obtained by the 512-dimensional scVI representation. The full scUNVEIL embedding exceeded this baseline by 8.7 macro-F1 points, despite scVI being optimized using the Hao et al. cells themselves.

Among competing zero-shot foundation-model representations, scFoundation was the strongest alternative, reaching 70.1 ± 2.0% macro F1. scUNVEIL improved on this result by 10.5 macro-F1 points. The zero-block Linear Embeddings ablation also reached 76.8 ± 1.8% macro F1, outperforming every evaluated non-scUNVEIL method and indicating that much of the gain arises from the measurement-enrichment objective even with no nonlinear residual blocks.

#### Enrichment outperforms PCA on the same training corpus

To test whether the gain exceeds conventional covariance-based compression, we fitted a 2048-component PCA to centered 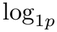 counts from the same 92.22 million training cells and 60,000 genes, without variance scaling. We used randomized SVD of the full gene covariance matrix and applied the frozen projection to Hao under the same classifier protocol. Pretraining PCA reached 69.1±2.5% macro F1, compared with 76.8±1.8% for linear scUNVEIL. This supports a benefit of measurement enrichment beyond preserving the highest-variance directions of the pretraining data.

#### Higher annotation performance is accompanied by lower inference cost

scUNVEIL generated embeddings for the complete benchmark in 00:01:41 on a single CPU. The corresponding runtime was 82:46:40 for scFoundation, 20:13:01 for scGPT, and 128:58:20 for the 33-layer UCE model. Consequently, the representation with the highest annotation performance was also orders of magnitude faster to generate than the strongest transformer-based alternatives. Although dedicated GPUs can further reduce inference time, we report CPU runtimes to demonstrate that scUNVEIL remains practical on commodity hardware without a dedicated accelerator.

#### Performance is retained under moderate dimensionality reduction

The scUNVEIL representation remained highly informative after PCA compression: the 512- and 128-dimensional variants outperformed every competing representation at its best evaluated dimensionality (Table 1), achieving 79.3 ±1.9% and 77.1 ±2.2% macro F1, respectively. The 50-dimensional variant reached 70.9±2.5% macro F1, below 71.9 ±2.3% for scVI with 512 dimensions.

Performance declined more substantially when the representation was compressed to 10 dimensions, where macro F1 reached 39.9 ±2.5%. One plausible explanation is that the PCA projection was fitted to embeddings from a broad human-cell atlas. Its leading components may therefore prioritize tissue- and organ-level variation rather than the comparatively subtle variation that distinguishes closely related PBMC subtypes. The largest performance decline occurred in this very-low-dimensional regime.

#### scUNVEIL remains effective at reduced input depth

We next evaluated annotation performance as the number of retained input UMIs was reduced (Figure 3). We evaluated seven retention probabilities from 0.1% to 100%, with 20 fresh thinning and donor-split repetitions per representation and retention. Counts were thinned before the 4/3/1 donor split, and the learning rates selected at full depth were reused. Interpolating the mean scUNVEIL curve linearly against log retention indicated approximately 88% fewer input UMIs to match scFoundation, the strongest competing foundation model, and approximately 94% fewer input UMIs to match full-depth raw 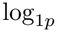 RNA measurements. These results indicate that the pretrained representation compensates for a substantial fraction of the information lost under shallow sequencing.

**Figure 1:**
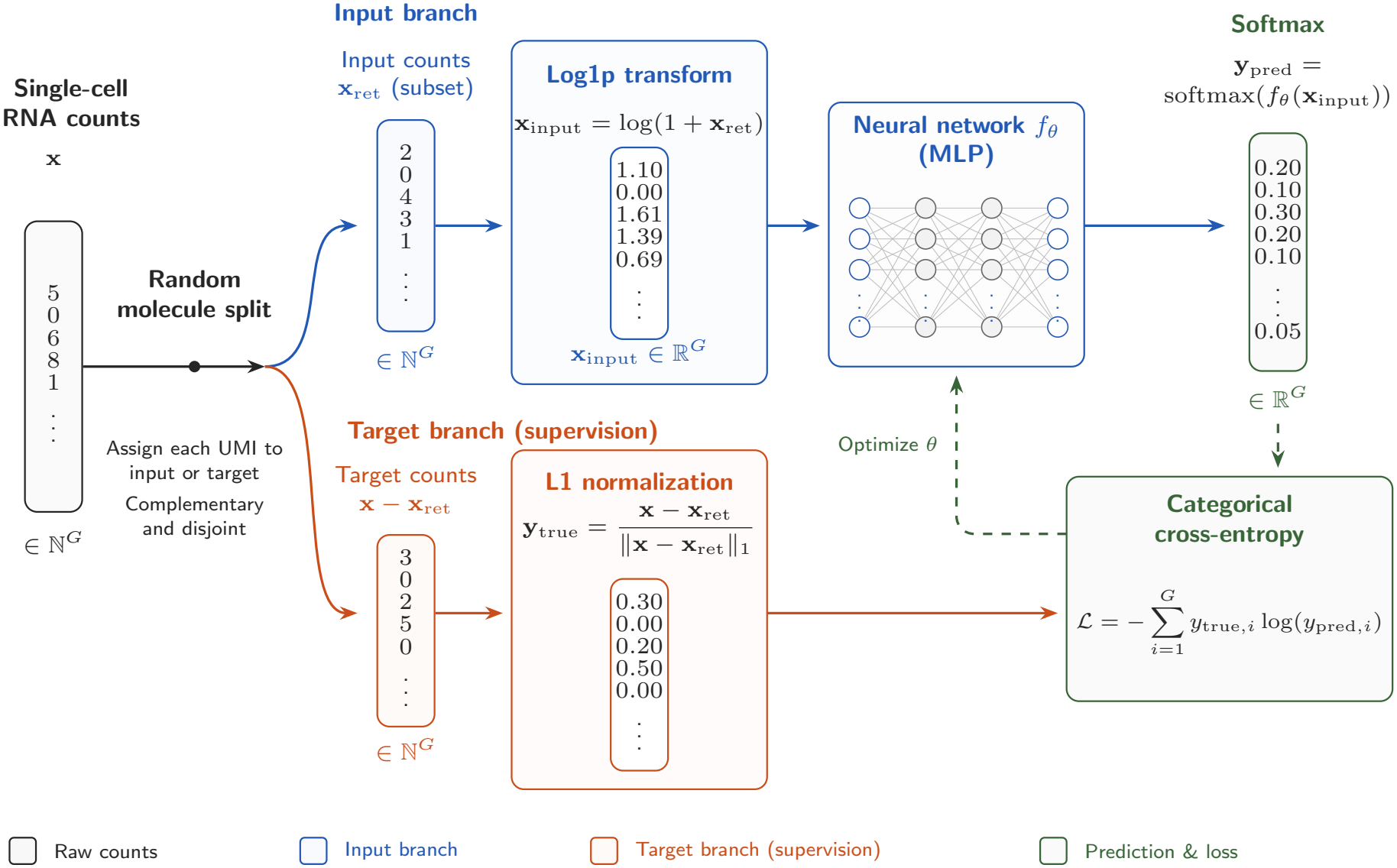
scUNVEIL pretraining objective. A random molecule split produces complementary input and held-out counts. Here **x** *∈* N^*G*^ is the observed count vector over *G* genes, **x**_ret_ contains retained input counts, and **x** − **x**_ret_ contains complementary held-out counts. N denotes nonnegative integers and R denotes real numbers. The log transform is applied elementwise, yielding the real-valued network input **x**_input_. The MLP *f*_*θ*_, with trainable parameters *θ*, produces gene logits; softmax converts them to predicted gene probabilities **y**_pred_. The target distribution **y**_true_ is obtained by dividing held-out counts by their L1 norm *∥*·*∥*_1_, the total held-out UMI count. *ℒ* is the categorical cross-entropy summed over gene index *i* = 1, …, *G*. Values and network widths are schematic.

**Figure 2:**
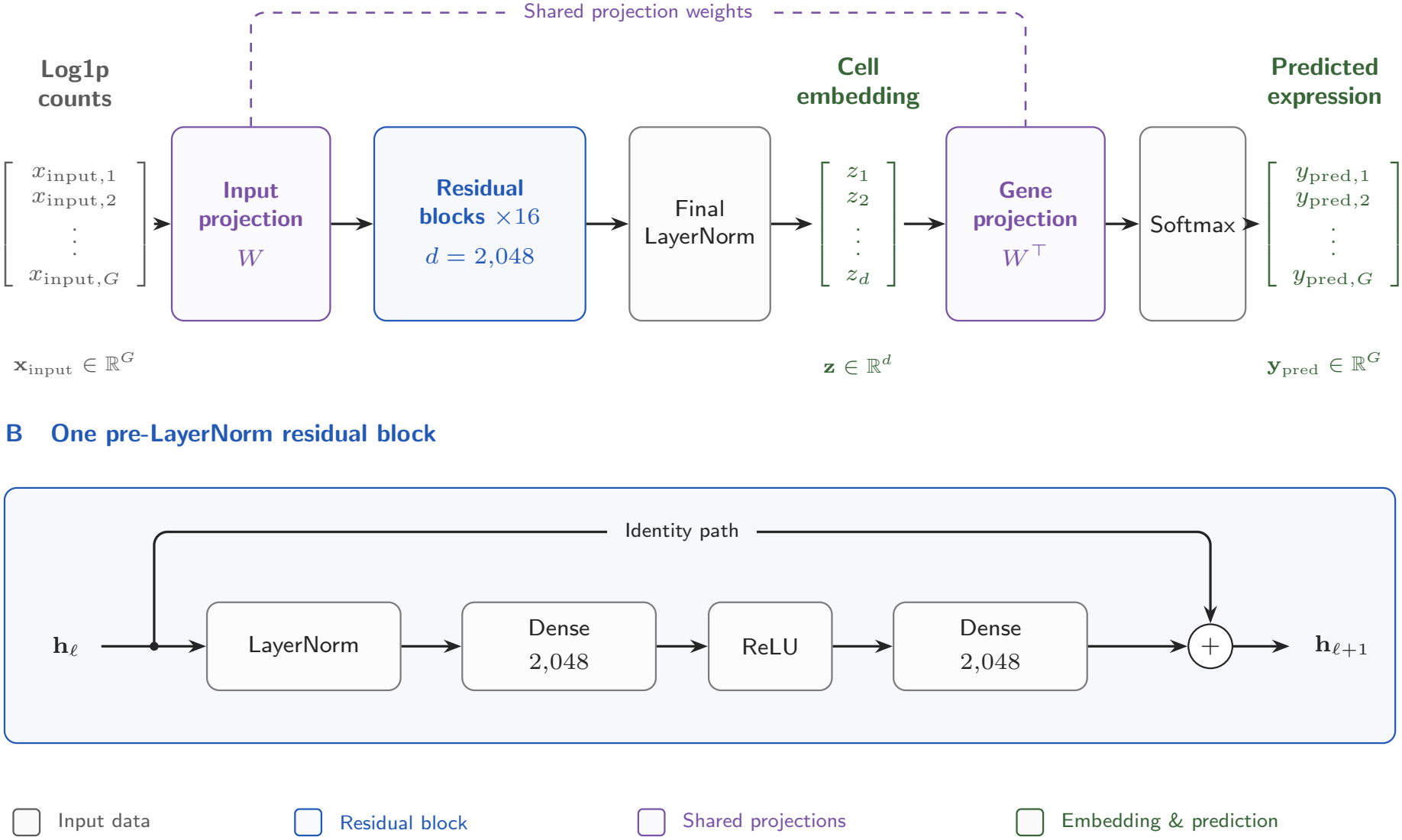
scUNVEIL model architecture. **(A)** Log-transformed counts **x**_input_ *∈* ℝ^*G*^ are projected into a *d*-dimensional space, processed by 16 sequential residual blocks, and normalized by a final LayerNorm to obtain the cell embedding **z** *∈* ℝ^*d*^. Here *G* = 60,000 is the gene vocabulary size and *d* = 2,048 is the embedding and hidden-layer width. The bias-free input projection uses *W ∈* R^*d*×*G*^; the gene projection reuses *W*^*⊤*^ to produce logits. Softmax converts these logits to the predicted gene probabilities **y**_pred_. Predicted expression can be reported as log_10_-CPM values log 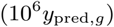 for gene *g*. The dashed link indicates weight sharing. **(B)** Within each residual block, the hidden state **h**_*ℓ*_ passes through LayerNorm, a dense layer of width 2,048, ReLU, and a second dense layer of width 2,048. An identity path bypasses this entire sequence and is added to the residual branch to produce **h**_*ℓ*+1_, where *ℓ* indexes the blocks.

**Figure 3:**
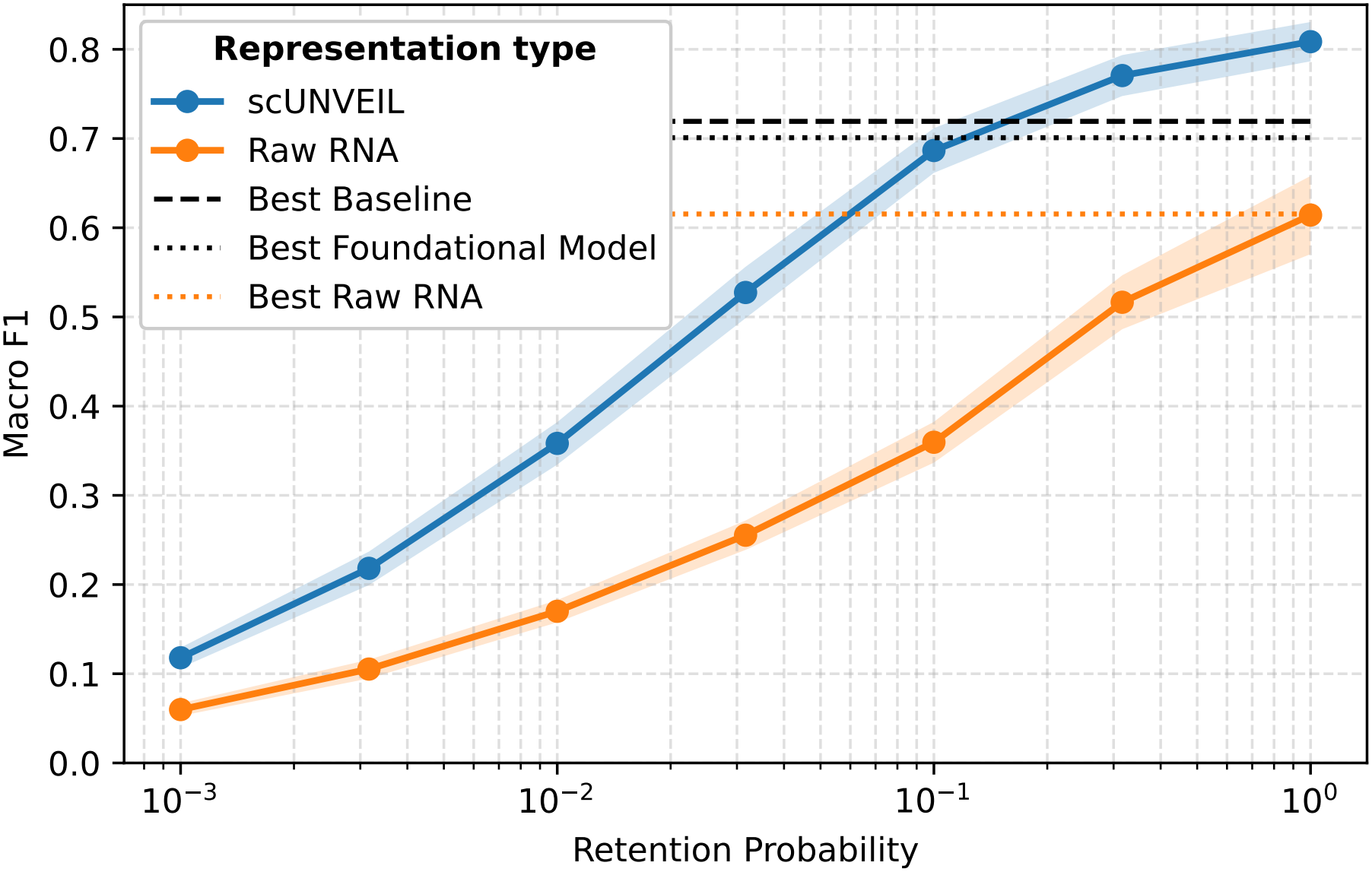
Macro F1 as a function of retained input UMI depth. Curves and shaded bands show mean ± sample SD over 20 random donor-level train–validation–test splits per representation and retention probability. Horizontal lines show full-depth means from the 100-repetition main benchmark. The interpolated scUNVEIL curve matches scFoundation with approximately 88% fewer input UMIs and full-depth raw 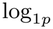 RNA with approximately 94% fewer.

### 4.2 Measurement-Enriched Expression Estimates

In addition to producing cell embeddings, scUNVEIL predicts a probability distribution over the complete gene vocabulary in the same forward pass. This distribution estimates the cell’s effective measured transcriptome composition from its finite UMI observation. For visualization and downstream analysis, we express the predicted probabilities on a log-counts-per-million scale as 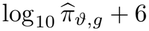.

We next evaluated whether these measurement-enriched gene-expression estimates preserve biologically useful gene-level information.

#### Measurement-enriched marker expression improves downstream annotation

We evaluated the predicted expression values through a downstream task with independently defined labels. Using the same 50 manually selected PBMC marker genes, we compared linear classification based on raw 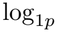-transformed counts, MAGIC-imputed expression, scVI-imputed expression, and scUNVEIL predictions.

The scUNVEIL marker-gene representation achieved 78.8 ±2.1% macro F1, compared with 70.1 ±2.3% for MAGIC and 57.2 ±2.3% for scVI (Table 1). The improvement over MAGIC was 8.7 macro-F1 points and was substantially larger than the variability across donor splits.

This comparison is particularly notable because MAGIC uses the complete Hao et al. dataset to construct its cell–cell smoothing structure, whereas scUNVEIL predicts the expression composition of each cell independently from that cell’s raw UMI counts. Despite using only 50 predicted marker values, the scUNVEIL representation also outperformed every competing foundation-model embedding evaluated in Table 1.

#### Qualitative marker analysis

Figure 4 compares population-level expression distributions for two representative marker-gene contrasts. For the canonical T-cell marker CD4, both MAGIC and scUNVEIL produce clear separation between CD4^+^ and CD8^+^ T-cell populations. For FOXP3, scUNVEIL produces stronger separation between regulatory T cells and CD4 central-memory T cells, illustrating its ability to recover informative structure for a rarer and more closely related cell population.

**Figure 4:**
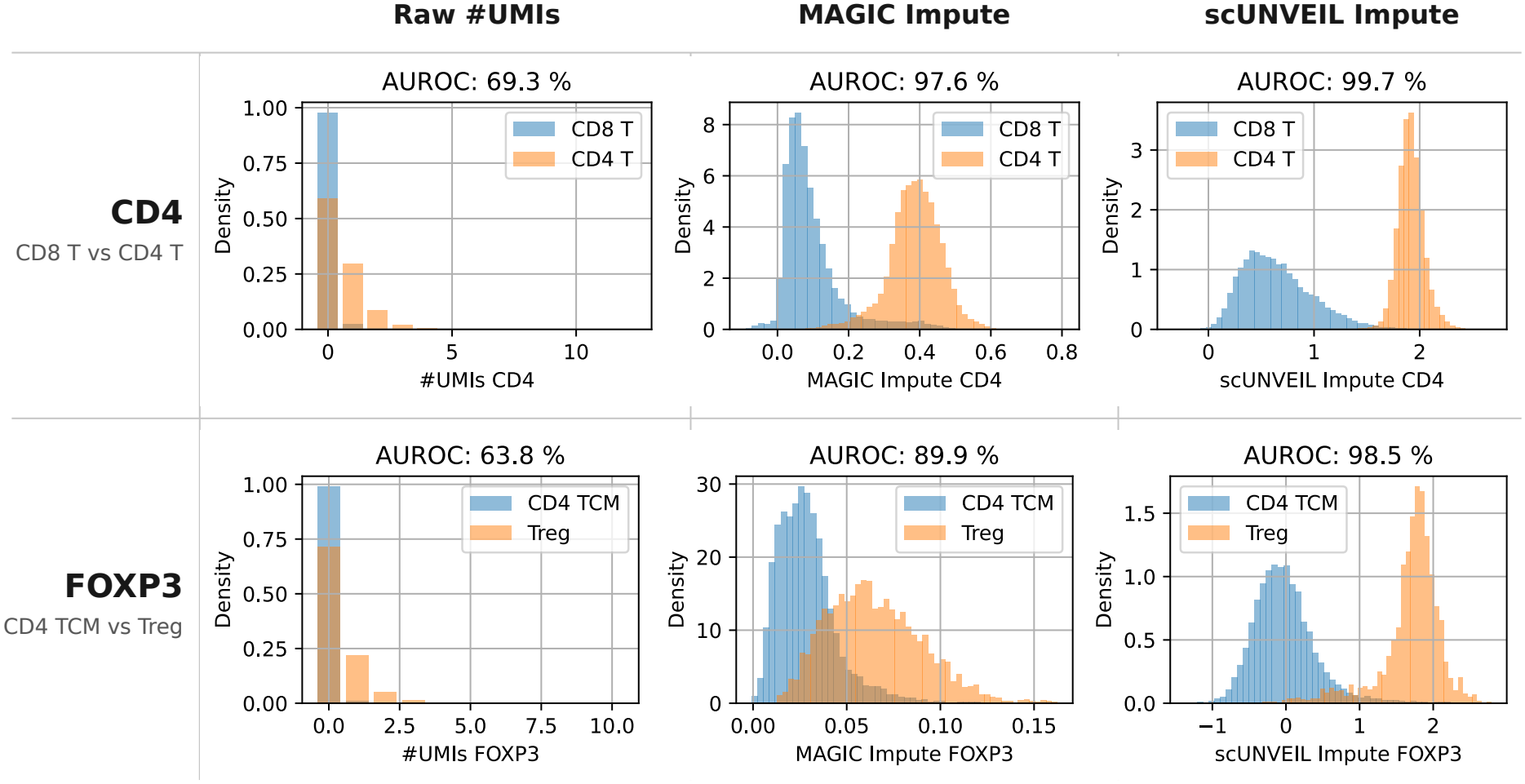
Population-level expression distributions for representative marker-gene contrasts. The top row compares CD4 expression between CD4^+^ and CD8^+^ T cells, and the bottom row compares FOXP3 expression between regulatory T cells and CD4 central-memory T cells. Columns show raw measured UMI counts, MAGIC-imputed expression, and scUNVEIL measurement-enriched expression in log_10_(CPM), respectively. The AUROC displayed above each panel quantifies separation between the two cell populations.

#### Interpretation and intended use

Together, the quantitative and qualitative results show that the measurement-enriched output contains gene-level signal that is useful for distinguishing closely related cell populations. They do not establish that the predicted value of every gene is an exact reconstruction of an unobserved cell-specific expression state. In particular, scUNVEIL estimates the effective measured transcriptome composition under the pretraining population and measurement process; it does not recover transcripts that are neither observed nor predictable from population-level regularities.

We therefore view the predicted expression values primarily as a tool for hypothesis generation, including the identification of candidate markers, cell populations, and contrasts for subsequent investigation. They should not be treated as confirmatory evidence for differential expression without validation using appropriate measurements and statistical analyses.

## 5 Supplementary experiments beyond cell-type annotation

Cell-type annotation provides a standardized comparison of representations. To test broader utility for single-cell phenotyping, we evaluated four prediction tasks with reference outcomes obtained through lineage tracing, functional assays, microscopy or fluorescence sorting (Table SE1). These targets were measured independently of RNA-based annotation. We tested whether frozen scUNVEIL embeddings improve prediction, particularly when RNA measurements become sparse through molecule dilution.

**Table SE1:** Downstream tasks with externally measured reference targets.

| Dataset | Measured target | Held-out unit | Metric |
|---|---|---|---|
| Pancreatic T cells | TCR recognition of autologous tumor cells | Patients | AUROC |
| Mitochondrial potential | TMRE-high versus TMRE-low T cells | Subjects | AUROC |
| SCOPE-seq2 | Cell area and circularity | Device regions (one patient) | *R*<sup>2</sup> |
| ReSisTrace | Lineage recovery after treatment or untreated growth | Sister-cell/lineage groups | AUROC |

### Shared evaluation protocol

We compared all 2,048 frozen scUNVEIL features with raw log(1 + *X*^*′*^) RNA and PCA of log1p RNA; the RNA baselines used all available gene features. PCA was fitted once on *all eligible cells at their original UMI depth*, including calibration and held-out cells, without using outcome labels. Its frozen gene means and loadings then projected the current sampled counts. PCA therefore has privileged access to the full-depth measurements of the very test cells later evaluated after dilution. We deliberately grant PCA this information advantage to make the comparison more demanding for scUNVEIL, whose frozen encoder processes each cell using only its current sampled RNA. We retained 2,048 PCs, capped at 955 for the 956-cell SCOPE-seq2 cohort. No library-size normalization, feature standardization or PCA whitening was applied.

Each repetition used a fresh grouped train/development/test split, fresh binomial thinning 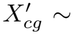 Binomial(*X*_cg_, *r*) at *r ∈* {1, 0.1, 0.01}, and freshly fitted linear predictors. Thinning affected all three roles; the 100% condition used original counts. Partitions and selected cells were shared across retention rates, and sampled counts were paired across representations. Eligibility and outcome labels remained fixed, including cells with no retained UMIs. Classification used L2-regularized logistic regression; morphology used ridge regression.

Regularization was selected once per representation, retention rate and task using reserved calibration groups, then fixed across repetitions. Calibration maximized the reported AUROC or region-level *R*^2^; morphology targets had separate ridge penalties. Logarithmic penalty grids were extended if an endpoint still improved the calibration objective. Calibration groups were excluded from subsequent repetitions, and development predictions did not retune or refit the models. All figures summarize the first 100 completed paired repetitions as mean ± sample SD. These overlapping resamples measure within-dataset variability conditional on fixed calibration and pretrained weights; they are not independent biological replicates or population confidence intervals. Retention is relative to each dataset’s original measurements, not a common absolute sequencing depth. Axis limits follow the displayed means and SDs; reference lines at AUROC = 0.5 or *R*^2^ = 0 appear only when within that range.

**Figure SE1:**
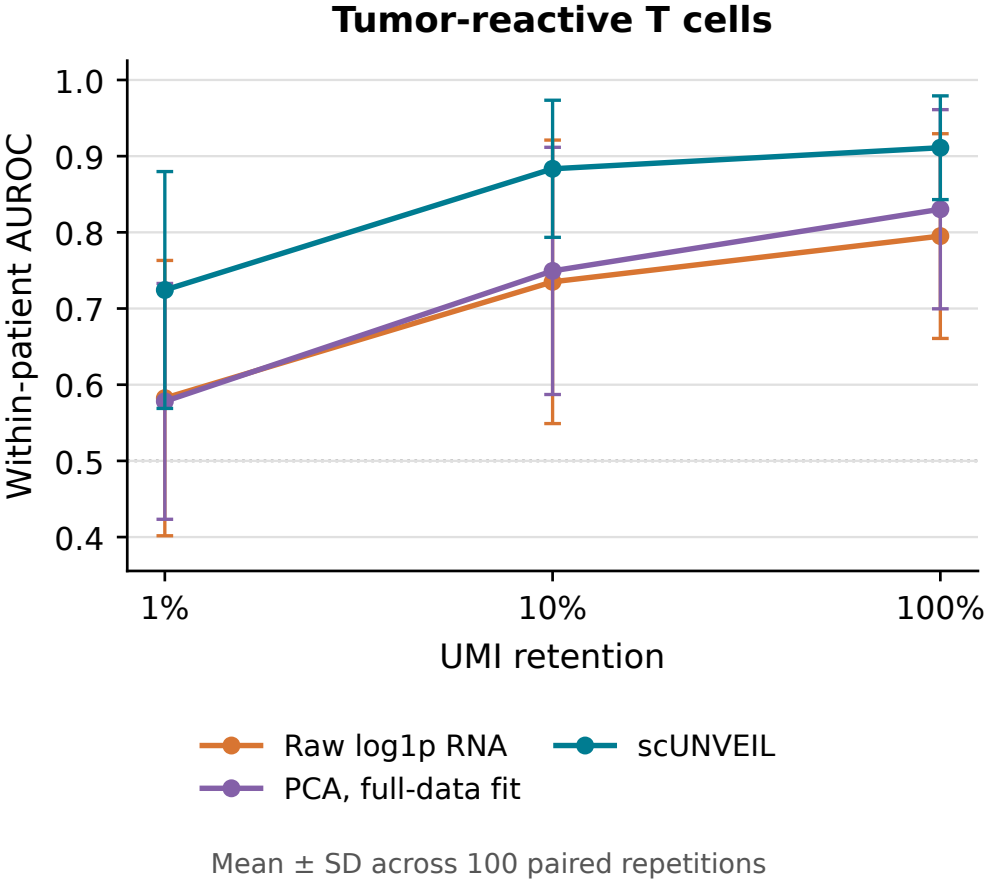
Tumor-reactivity prediction. Mean within-patient test AUROC ± sample SD across 100 paired patient-disjoint repetitions, using one cell per tested clonotype.

#### 5.1 Tumor-reactive T cells

The source study functionally tested T-cell receptors (TCRs) for recognition of tumor cells from the same patient (Meng et al., 2023; [GSE254250 study](https://doi.org/10.1126/scitranslmed.adh9562)). The prediction target is binary: whether the sequenced cell’s TCR clonotype was experimentally classified as tumor-reactive or nonreactive. The task is to infer this assayed recognition phenotype from a single cell’s RNA alone.

We used 118 tested clonotypes from nine pancreatic-cancer patients, linked to 3,990 RNA profiles. Two patients were reserved for calibration, using leave-one-patient-out validation. Each repetition assigned the remaining seven patients to four training, one development and two test patients, with one freshly selected cell per clonotype and class-balanced training. AUROC was averaged within mixed-label test patients; five patients supplied this primary test evidence. Patient separation did not fully separate sequencing libraries.

At 10% retention, mean AUROC was 0.883 for scUNVEIL, 0.749 for PCA and 0.735 for raw RNA (Figure SE1). scUNVEIL also exceeded both RNA baselines at full depth and 1% retention. Labels concern selected, assayed receptors rather than direct killing by each sequenced cell or clinical response.

#### 5.2 Mitochondrial-potential phenotype

The source study stained peripheral-blood T cells with tetramethylrhodamine ethyl ester (TMRE), a potential-sensitive fluorescent dye, and used fluorescence-activated cell sorting to separate high- and low-fluorescence populations before RNA sequencing (Lindau et al., 2026; [GSE315510 study](https://doi.org/10.1093/jimmun/vkag120)). The binary prediction target is the experimentally measured TMRE-high versus TMRE-low gate. The task is to infer this mitochondrial-potential-associated fluorescence phenotype from RNA alone.

We used five lung-cancer subjects with two visits each; original-count QC retained 73,180 cells. One subject was reserved for cross-visit calibration; each repetition split the remaining four subjects into two training, one development and one test subject. Both visits and gates remained together. We sampled at most 250 cells per gate/visit, balanced the gates, and weighted subject/visit/gate combinations equally in training. Test AUROC was calculated within each visit and averaged over the test subject’s visits.

Mean scUNVEIL AUROC exceeded both RNA baselines at every retention (Figure SE2). At full depth it reached 0.910 versus 0.887 for PCA; at 1% the corresponding values were 0.719 and 0.597. High/low gates were sequenced in separate libraries, leaving potential systematic processing effects. TMRE gate membership is a potential-associated phenotype, not a direct measure of ATP production or metabolic flux.

**Figure SE2:**
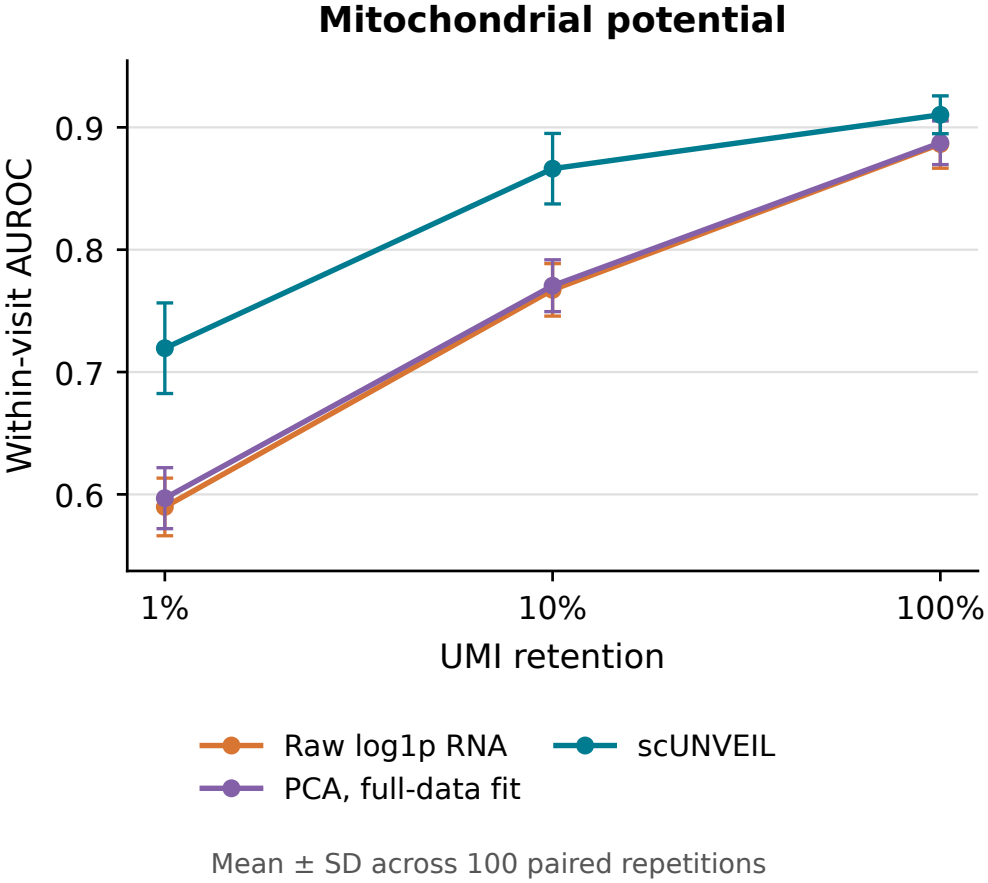
Mitochondrial-potential phenotype prediction. Within-visit test AUROC for TMRE-high versus TMRE-low T cells, mean ± sample SD across 100 paired subject-disjoint repetitions. Four non-calibration subjects supply the evaluation evidence.

#### 5.3 Cell size and shape

The SCOPE-seq2 study imaged individual live cells and linked their microscopy measurements to RNA profiles from the same cells (Liu et al., 2020; [GSE151137 study](https://doi.org/10.1038/s41598-020-76599-w)). We predict two continuous imaging measurements separately from RNA alone: cell area, describing size, and circularity, describing how round the cell appears. Targets are the authors’ released standardized imaging values, not quantities inferred from RNA.

We used 956 cells from one glioblastoma specimen. Two of ten technical device regions were reserved for leave-one-region-out calibration; the remaining eight were split into five training, one development and two test regions per repetition. *R*^2^ was calculated within each test region and averaged equally. These regions assess transfer within one specimen, not across patients.

At 10% retention, cell-area *R*^2^ was 0.208 for scUNVEIL, 0.056 for raw RNA and 0.018 for PCA (Figure SE3). For circularity, PCA was more competitive: 0.125 for scUNVEIL versus 0.107 for PCA. At 1%, scUNVEIL’s mean *R*^2^ was only 0.039 for area and 0.038 for circularity; PCA approximately matched the area result. Absolute predictive performance was therefore limited at the most severe dilution, despite improvements over raw RNA.

**Figure SE3:**
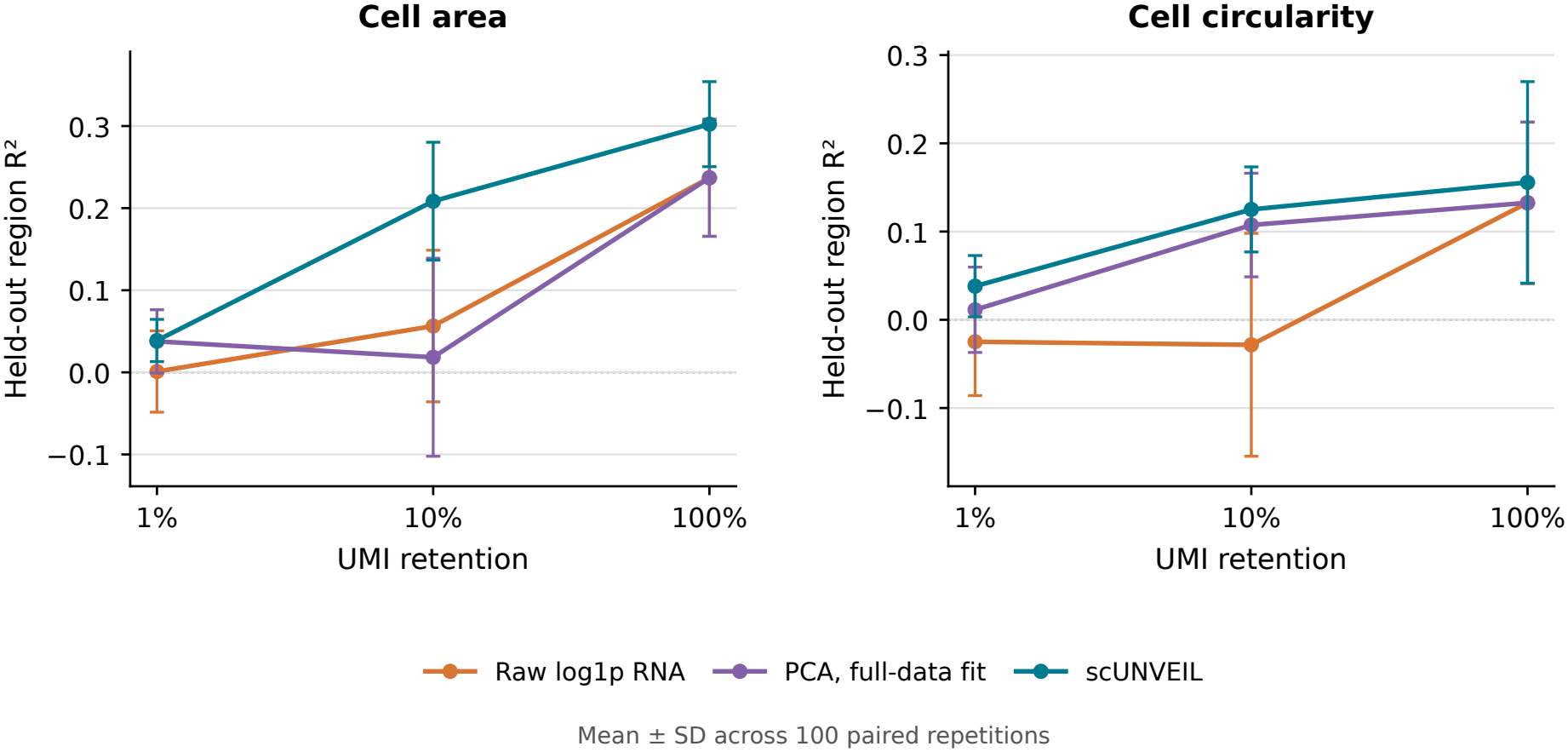
Microscopy-derived cell size and shape. Mean held-out-region *R*^2^± sample SD across 100 paired repetitions. Full-depth raw-RNA and PCA curves overlap: the 955-PC basis spans the observed cell variation. Negative scores are retained.

#### 5.4 Lineage recovery after treatment

ReSisTrace [7] used sister-cell tracing to link pretreatment ovarian-cancer transcriptomes to subsequent recovery of related lineages after carboplatin, olaparib, NK-cell exposure or untreated growth. For each condition, the binary target is whether a lineage was recovered afterward or not detected. The task is to predict that later lineage outcome from pretreatment RNA, using the study’s released recovery annotations.

We used 32,642 annotated cells. Connected sister-cell and family identifiers defined indivisible groups within each source sample; 20% were reserved for calibration, and the remainder was split 60/20/20% within condition and biological replicate. Thus both source replicates contribute to training and testing. Calibration used three grouped folds. Training weights balanced outcome classes and lineage contributions; direct cell probabilities were averaged within test groups before AUROC calculation.

scUNVEIL improved mean AUROC over raw RNA at 10% and 1% retention in all four conditions, but PCA narrowed the advantage (Figure SE4). For carboplatin at 1%, mean AUROC was 0.668 for scUNVEIL, 0.648 for PCA and 0.621 for raw RNA. At 10%, olaparib scores were 0.742, 0.716 and 0.701, respectively. At 1%, PCA approximately matched scUNVEIL in the other three conditions; it exceeded scUNVEIL at full depth for carboplatin and untreated recovery. Recovery can reflect growth and sampling as well as resistance, so these results concern lineage-associated outcomes rather than treatment-specific resistance alone.

**Figure SE4:**
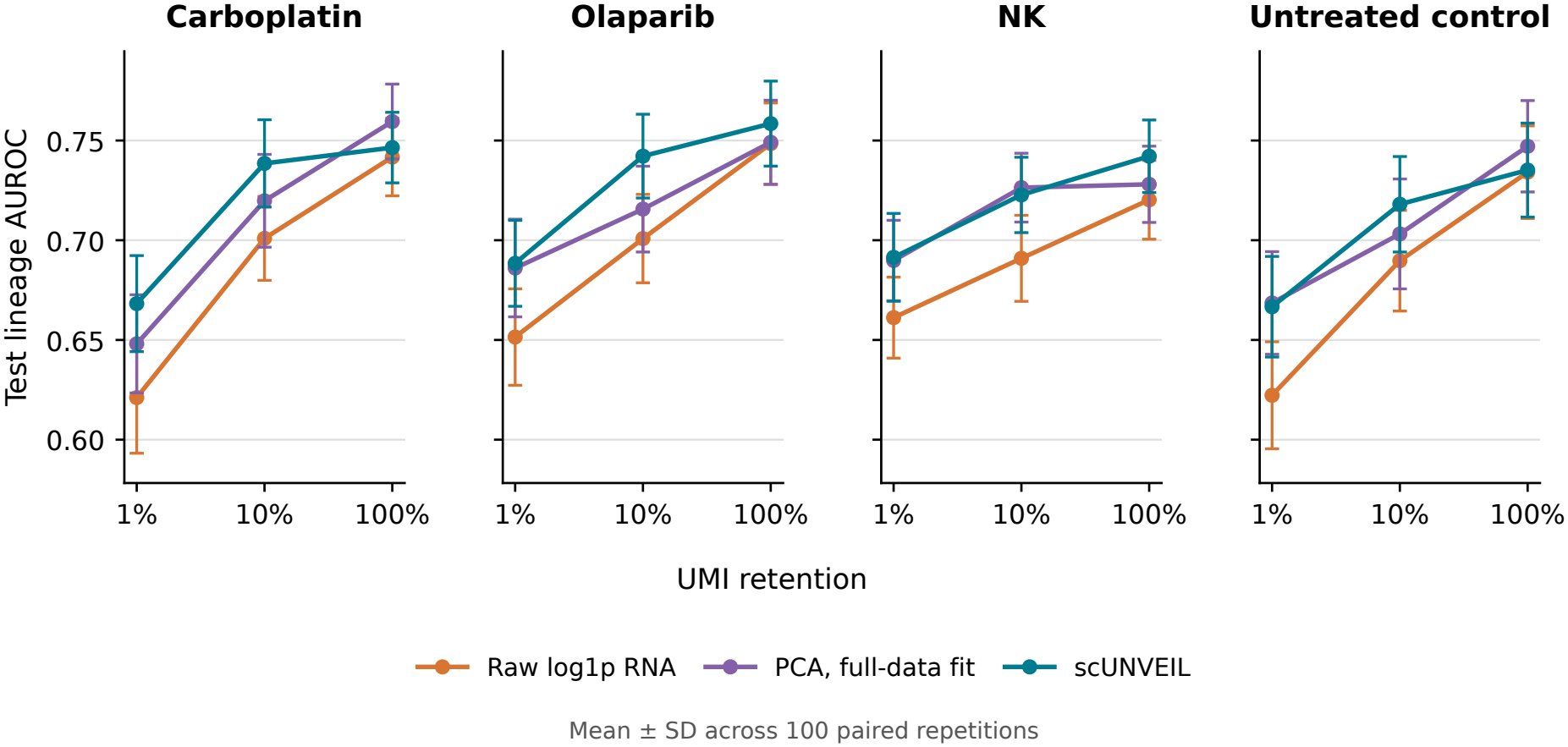
Lineage-recovery prediction. Test lineage AUROC at three UMI retentions, mean ± sample SD across 100 paired repetitions. PCA uses a frozen full-data, full-depth fit.

### Interpretation

These experiments support predictive utility beyond cell-type annotation, with particularly clear advantages over raw RNA and full-data PCA for tumor reactivity and the mitochondrial-potential phenotype. Morphology and ReSisTrace show more selective gains once PCA is included. Molecular thinning exposes useful transfer regimes, but advantages depend on the task and retained depth. Improved prediction from diluted RNA suggests that pretrained representations may reduce the sequencing depth needed for some downstream phenotyping tasks. Prospective shallow-sequencing studies could establish the achievable savings. The tasks were selected after exploratory screening and do not estimate performance across unselected phenotypes. Comparisons with other pretrained representations and independent validation cohorts would further establish the generality of these findings.

## 6 Discussion

scUNVEIL demonstrates that measurement enrichment can produce strong zero-shot representations using a simple single-vector MLP architecture. On fine-grained PBMC annotation, its embeddings outperformed the evaluated single-cell foundation models while requiring substantially less inference compute, and its measurement-enriched expression estimates were highly informative using only a small set of marker genes. The main methodological contribution is the next-UMI formulation of measurement enrichment and the representation-learning framework that follows from it. This formulation identifies the Bayes-optimal next-UMI predictor with the posterior mean of the cell’s measured composition in the infinite-depth limit. Because UMI order is irrelevant to this objective, the cell can be processed directly as a count vector, while the shared hidden state is implicitly trained to retain the information needed to predict additional UMIs. These results do not imply that transformers are generally unsuitable for scRNA-seq, but they show that attention over gene tokens is not necessary for obtaining strong representations in this setting.

The additional experiments extend this evidence beyond cell-type annotation to predicting tumor reactivity established through functional assays, mitochondrial potential assessed by fluorescence sorting, cell size and shape measured by microscopy, and subsequent lineage recovery tracked through lineage tracing. These predictions could help identify tumor-reactive T cells, characterize cellular physiology and morphology, and anticipate which lineages are likely to recover after treatment or untreated growth. Frozen scUNVEIL embeddings showed particularly clear advantages over raw RNA and PCA for tumor reactivity and mitochondrial potential, while morphology and lineage recovery showed more selective gains depending on the task and retained depth. These findings support the broader motivation for measurement enrichment: population-level pretraining can make sparse RNA measurements more informative for predicting independently observed properties of cells.

The broadest comparison with competing representations remains the Hao et al. PBMC benchmark. The additional phenotype experiments extend the range of evaluated tasks, while further datasets and pretrained-model comparisons would help establish how broadly these advantages transfer. Gene-expression estimates were evaluated primarily through their utility for PBMC annotation. The interpretation of the model should also remain tied to its training objective: scUNVEIL estimates the effective measured transcriptome composition that would be observed at greater UMI depth under the same measurement process. It does not attempt to remove systematic technical effects, recover a distortion-free biological transcriptome, or perform batch correction. Similarly, its predicted expression values are most appropriately treated as estimates for exploratory analysis and hypothesis generation rather than as conclusive evidence of differential expression.

scUNVEIL is straightforward to deploy: a single pip install scunveil command installs the package, pretrained weights are downloaded automatically, and a few lines of Python generate embeddings or enriched expression estimates. In our benchmark, it processed thousands of cells per second on a consumer CPU.

The computational efficiency of scUNVEIL may also create opportunities for further scaling. Additional compute could be directed toward larger models, longer training, or broader pretraining corpora. Whether such scaling would produce further gains remains to be tested, but the current model is unlikely to represent the practical limit of the approach.

Furthermore, measurement enrichment provides a concrete way to reason about the limits of pretraining. For a proposed downstream task, one can ask whether the task would become easier if the same cells were measured at substantially greater UMI depth under the same effective observation process. If so, pretraining may help by estimating information that is absent from the sparse measurement but predictable from population-level regularities. If even an arbitrarily deep measurement would not simplify the task, measurement enrichment alone provides little reason to expect a substantial improvement. This perspective does not claim that measurement enrichment is the only possible source of useful representations, but it offers a clear reference point for connecting a pretraining objective to the downstream information it can plausibly recover.

## Supporting information

Supplementary Table S1: Pretraining datasets

## Code and Weights Availability

The scUNVEIL project is fully open source and released under the Apache License 2.0. The main `scunvei`l package provides inference utilities for generating cell embeddings and measurement-enriched gene-expression estimates from raw UMI count vectors. The package is distributed through GitHub and PyPI. When the package is installed via `pip install scunveil`, the required pretrained model weights and fitted PCA projection are downloaded automatically from Hugging Face during use. Complete training and experimental reproducibility code is also publicly available.

**Inference package (**v0.6.0)

GitHub: https://github.com/thonzyk/scunveil

PyPI: https://pypi.org/project/scunveil/

Hugging Face: https://huggingface.co/thonzik/sc-unveil

**Training pipeline (**v0.6.0)

GitHub: https://github.com/thonzyk/scunveil-training

**Experimental reproducibility (**v0.6.0)

GitHub: https://github.com/thonzyk/scunveil-experiments

## AI Use Statement

Large Language Models were used to assist with rephrasing, structuring sections of the text, identifying relevant literature, and harmonizing mathematical notation. All content was reviewed, verified, and finalized by the authors.

## Acknowledgements

This work was supported by ELIXIR CZ Research Infrastructure (ID LM2023055, MEYS CR). Computational resources were provided by the e-INFRA CZ project (ID 90254), supported by the Ministry of Education, Youth and Sports of the Czech Republic.

## Supplementary Information

## Appendix A: Mathematical derivation of measurement enrichment

This appendix derives the statistical target of the next-UMI prediction objective introduced in Section 3.3. Unless stated otherwise, expectations are taken under the valid-split training distribution, conditioned on *N*^out^ *>* 0.

### 6.1 High-depth counterfactual

For a hypothetical measurement of the same effective composition ***π***_c_ at depth *K*, define

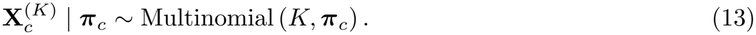

By the strong law of large numbers,

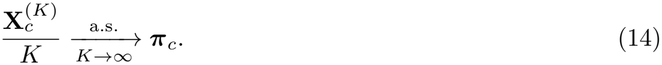

Thus, ***π***_c_ is the normalized high-depth limit of the measurement under the same effective observation channel. This is a mathematical idealization rather than a claim that infinitely many unique molecules can be obtained from a finite cell.

The multinomial model in Equation (3) can be interpreted as an approximation to sampling from a large assay-accessible molecular population. It also arises exactly when independent Poisson-distributed gene counts are conditioned on their total count.

### 6.2 Single-UMI prediction

Consider the conceptual task of predicting one UMI *U*^out^ selected uniformly from the held-out set. Given an input count vector **x**, define the single-UMI cross-entropy loss as

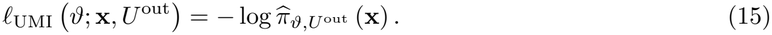

Conditional on the cell-specific effective measured composition,

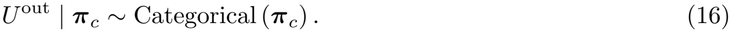

Therefore, for a fixed input realization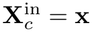

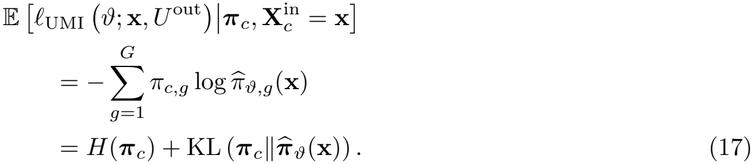

Because *H*(***π***_c_) does not depend on the model parameters, minimizing the expected loss for a cell with known ***π***_c_ is equivalent to minimizing the divergence between the cell’s effective measured composition and the predicted composition.

Equation (17) describes the oracle target for a particular cell. In practice, however, the model observes only the sparse input **x**, and multiple cell-specific compositions may be compatible with that input.

### 6.3 Population risk and the Bayes-optimal predictor

For a fixed input vector **x**, define

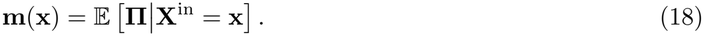

For any candidate prediction **q** *∈* Δ^G*−*1^, its conditional population risk is

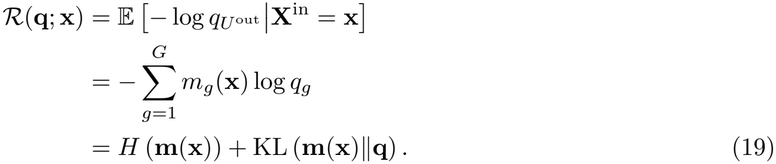

The conditional risk is therefore minimized by

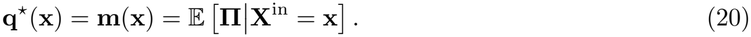

Equivalently,

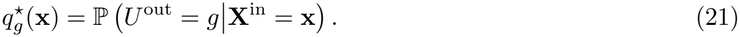

Thus, the Bayes-optimal next-UMI predictor is simultaneously the posterior predictive distribution of the next held-out UMI and the posterior mean of the effective measured composition. This establishes Equation (9).

The distinction between ***π***_c_ and **q**^*⋆*^(**x**) is important. The former is the effective measured composition of a particular cell. The latter is the optimal estimate of that composition available from the sparse input **x** under the population distribution represented during pretraining.

For an empty input, the same result gives:

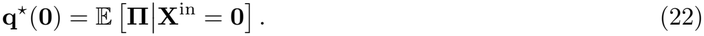

### 6.4 Empirical held-out objective

The implemented objective uses the empirical held-out composition from Equation (7):

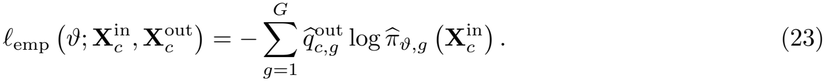

Let 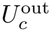 be sampled uniformly from the observed held-out UMIs. Conditional on the full split,

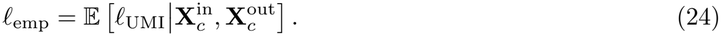

The empirical loss is therefore the Rao–Blackwellized form of the single-UMI loss with respect to the full held-out count vector.

Conditional on 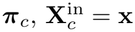, and 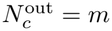,

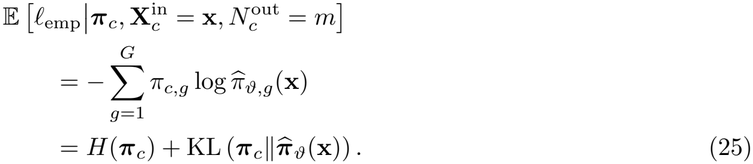

Hence, the single-UMI and empirical objectives have the same conditional expectation and the same expected gradient.

Under the multinomial idealization, the empirical loss is an average of 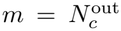 conditionally independent single-UMI losses. Its conditional variance is

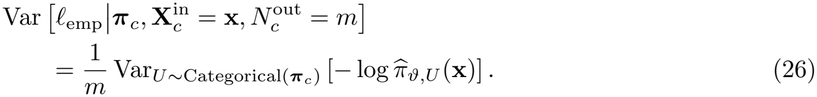

Using all held-out UMIs therefore reduces the conditional variance by a factor of *m* relative to using one independently sampled held-out UMI. The variance is equal when *m* = 1 and is smaller whenever *m >* 1 and the single-UMI loss is nonconstant.

Averaging over cells, compositions, input depths, and random splits gives

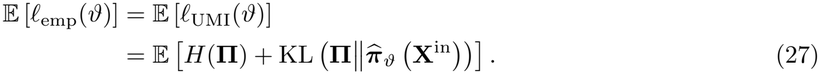

The implemented loss is therefore next-UMI prediction in expectation, while using the complete held-out count vector to obtain a lower-variance gradient estimate.

### 6.5 Randomization over input depth

Under the idealized procedure, the retention probability is sampled as *r ~* Uniform(0, 1), and each of the *N*_c_ observed UMIs is retained independently with probability *r*. Marginalizing over *r* gives

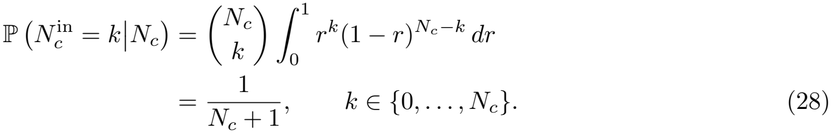

Thus, before excluding empty held-out sets, the induced input depth is uniform over all integer depths from 0 to 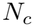. Conditioning on 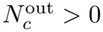 gives

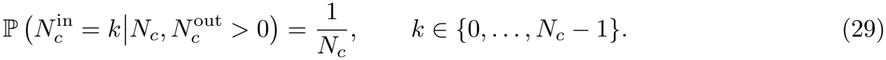

The idealized objective therefore weights every usable input depth equally within each cell.

In the implemented training procedure, truncating the retention distribution to

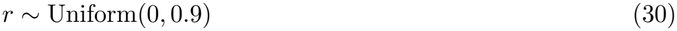

reweights the encountered input depths and reduces the frequency of nearly complete input sets. This changes the weighting of training examples but preserves the conditional task: predict the held-out UMI distribution from the retained counts.

The input depth is directly available from the count vector:

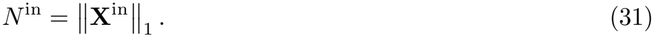

Consequently, scUNVEIL does not require a separate source-depth token and does not receive or predict a requested target depth. At every input depth, the output is a probability distribution intended to estimate the same depth-independent quantity, ***π***_c_.

### 6.6 Consistency in the high-depth limit

Let 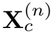 denote an observation of the effective composition ***π***_c_ at input depth *n*. By Equation (14),

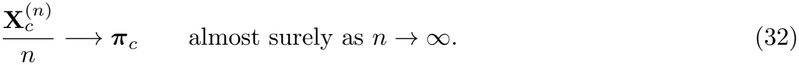

Under standard identifiability and posterior-consistency conditions, the posterior distribution over **Π** concentrates on ***π***_c_, and its posterior mean satisfies

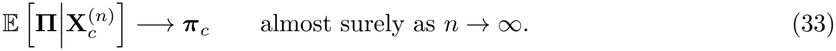

At finite depth, the Bayes-optimal estimator combines cell-specific evidence with population-level regularities. At increasing depth, the observed cell supplies progressively more information and the influence of the population prior diminishes.

Equation (33) is a property of the ideal Bayes predictor. A finite neural network trained on finite data approximates this predictor only to the extent permitted by the training distribution, model capacity, optimization procedure, and available data.

### 6.7 Interpretation of the output

The model output 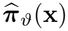 is the estimated composition itself, not a count vector at a user-specified depth. It has two equivalent interpretations:

1. The predicted distribution of an additional UMI generated under the same effective observation process.
2. The estimated normalized limit that would become apparent as the finite-sampling uncertainty of that process is reduced.

The log-counts-per-million transformation in Equation (12) expresses the predicted composition on a convenient scale but does not condition the prediction on a measurement depth of one million UMIs.

